# Unsupervised integration of multimodal dataset identifies novel signatures of health and disease

**DOI:** 10.1101/432641

**Authors:** Ilan Shomorony, Elizabeth T. Cirulli, Lei Huang, Lori A. Napier, Robyn R. Heister, Michael Hicks, Isaac V. Cohen, Hung-Chun Yu, Christine Leon Swisher, Natalie M. Schenker-Ahmed, Weizhong Li, Andrew M. Kahn, Timothy D. Spector, C. Thomas Caskey, J. Craig Venter, David S. Karow, Ewen F. Kirkness, Naisha Shah

**Affiliations:** Human Longevity, Inc., San Diego, CA 92121; Electrical and Computer Engineering, University of Illinois at Urbana-Champaign, IL 61820; J. Craig Venter Research Institute, San Diego, CA 92037; Division of Cardiovascular Medicine, University of California San Diego School of Medicine, La Jolla, California, 92093; Department of Twin Research and Genetic Epidemiology, King’s College London, U.K.; Molecular and Human Genetics, Baylor College of Medicine, Houston, TX 77030

## Abstract

Modern medicine is rapidly moving towards a data-driven paradigm based on comprehensive multimodal health assessments. We collected 1,385 data features from diverse modalities, including metabolome, microbiome, genetics and advanced imaging, from 1,253 individuals and from a longitudinal validation cohort of 1,083 individuals. We utilized an ensemble of unsupervised machine learning techniques to identify multimodal biomarker signatures of health and disease risk. In particular, our method identified a set of cardiometabolic biomarkers that goes beyond standard clinical biomarkers, which were used to cluster individuals into distinct health profiles. Cluster membership was a better predictor for diabetes than established clinical biomarkers such as glucose, insulin resistance, and BMI. The novel biomarkers in the diabetes signature included 1-stearoyl-2-dihomo-linolenoyl-GPC and 1-(1-enyl-palmitoyl)-2-oleoyl-GPC. Another metabolite, cinnamoylglycine, was identified as a potential biomarker for both gut microbiome health and lean mass percentage. We also identified an early disease signature for hypertension, and individuals at-risk for a poor metabolic health outcome. We found novel associations between an uremic toxin, p-cresol sulfate, and the abundance of the microbiome genera *Intestinimonas* and an unclassified genus in the *Erysipelotrichaceae* family. Our methodology and results demonstrate the potential of multimodal data integration, from the identification of novel biomarker signatures to a data-driven stratification of individuals into disease subtypes and stages -- an essential step towards personalized, preventative health risk assessment.

## Introduction

Despite the enormous U.S. healthcare spending of $3.3 trillion in 2016^1^, one in three individuals aged 50-74 years die prematurely from major age-related chronic diseases^2–4^. Challenging the status quo of reactive healthcare, preventative medicine offers an alternative means to better health for lower cost^5^. One approach to move beyond traditional medicine to more predictive, preventive practices is via systems medicine. As defined by Hood *et al*.^6^, systems medicine is the application of systems biology to the challenges of human health and disease. An interdisciplinary approach that measures, integrates, analyzes, and interprets a variety of clinical and non-clinical data is critical for a deeper understanding of the mechanisms that determine health and disease states. Significant computation and statistical analysis are essential to sift through large, diverse datasets and search for patterns, whether related to specified biological processes or to stratify complex diseases into distinct subtypes for health assessment.

Recent studies have shown the utility of collecting and analyzing diverse high-throughput data using unsupervised computational methods for more comprehensive insights into biological systems. Argelaguet *et al.*^7^ showed a need for such integrated analysis by introducing a computational framework of unsupervised integration of heterogeneous data and showed its utility by identifying major drivers of variation in chronic lymphocytic leukemia. Price *et al.*^8^ revealed communities of related analytes associated with diseases using unsupervised network analysis on a multimodal dataset.

In our previous work^4^, we introduced a platform of deep quantitative multimodal phenotyping that seeks to provide a comprehensive, predictive, preventative, and personalized assessment of an individual’s health status. The offered multimodal assays include whole genome sequencing, advanced imaging, metagenomic sequencing, metabolome, and clinical labs. This platform provides critical data not only to identify previously undiagnosed disease states but also to identify early disease biomarkers. Here, we present an analysis of the multimodal datasets that were collected for 1,253 self-assessed healthy adults and an independent validation dataset consisting of 1,083 adults with longitudinal data. To the best of our knowledge, this is the largest cohort with such a wide range of data modalities analyzed to date.

We built on the unsupervised approaches described above by performing a more comprehensive analysis to not only find novel patterns in disease risk but also to stratify individuals based on novel biomarker signatures to identify current disease states and early disease transition states. We performed an ensemble of machine learning analyses, including cross-modality associations, formation of modules of densely connected features to identify key biomarkers, clustering individuals into distinct health risk groups with corresponding biomarker signatures, and enrichment of longitudinal outcomes of individuals within each risk group. By doing so, we showcase a method to assess health status for preventative medicine by creating signatures of health and disease through multimodal data integration.

## Results

We collected data from 1,253 self-assessed healthy adults (median age 53; 63% male) across several modalities (Figure 1A), including whole-genome sequencing (WGS)^9^, microbiome^10^, global metabolome^11^, laboratory-developed tests for insulin resistance and prediabetes (Quantose™ ^12^), magnetic resonance imaging (MRI), computed tomography (CT) scan, routine lab work, vitals and personal/family medical history. Not all individuals were measured for all modalities (Table S1). For each of the modalities, several data features were measured, totaling 1,385 features from all modalities (see Methods). A majority of the cohort were of European ancestry (71.6%). The remainder were of East Asian (6.4%), Central/South Asian (3.4%), Middle Eastern (0.4%), African (0.3%), and admixed ancestries (18.0%).

**Figure 1.**
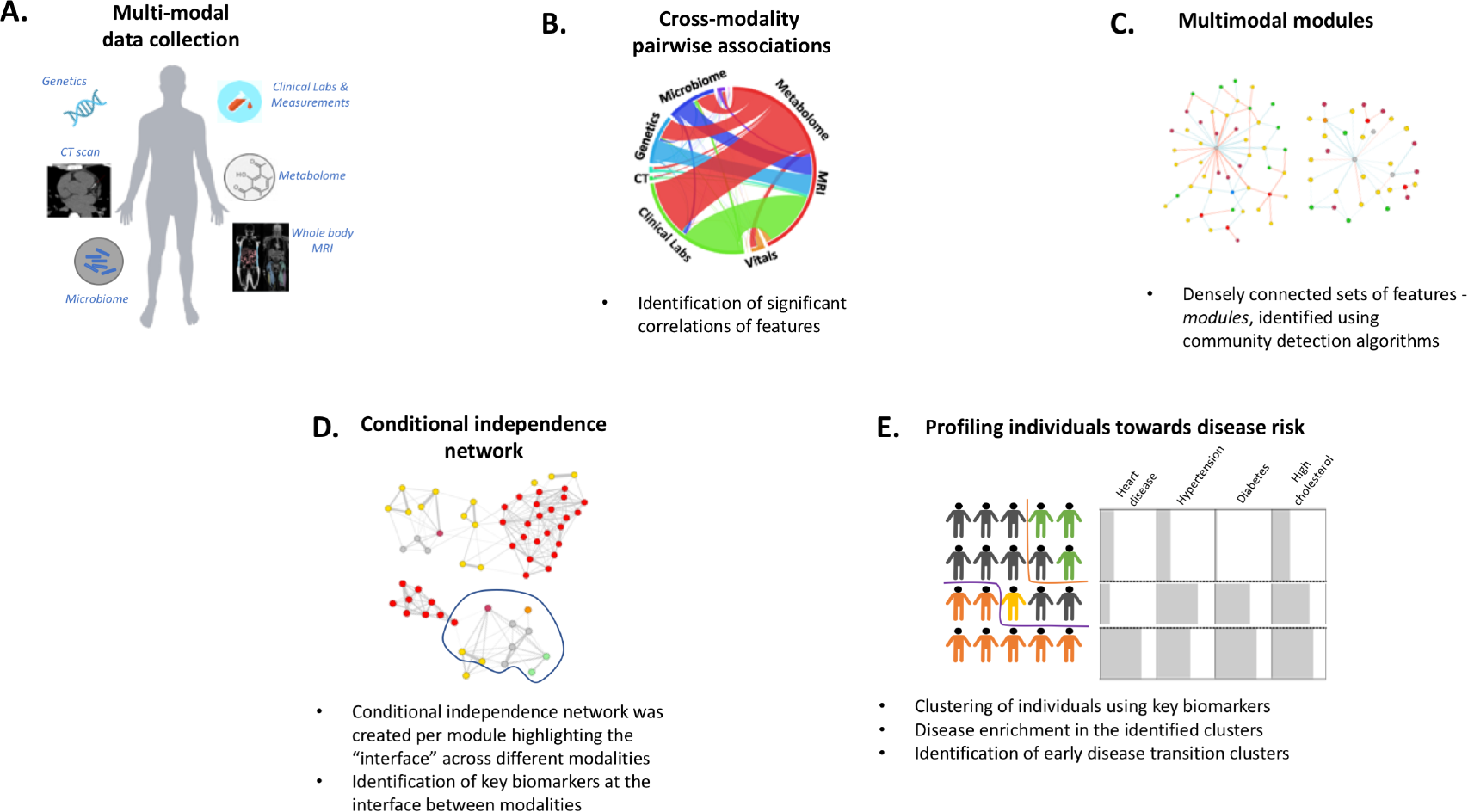
**(A)** In the study, we collected multi-modal data (n= 1,385 features) from 1,253 individuals. **(B)** We analyzed the data by performing cross-modality associations between features after correcting for age, sex and ancestry. **(C)** Using the associations, we performed community detection analysis and found modules of densely connected features. **(D)** To reduce the number of indirect associations and identify key biomarker features, we performed conditional independence network analysis (also referred to as a *Markov Network*). **(E)** Using the identified key biomarkers, we clustered individuals into distinct groups of health profiles. We characterize the clusters and perform disease risk enrichment analysis.

We performed four primary analyses using the collected multimodal data as summarized in Figure 1. First, we identified statistically significant associations across the data modalities (Figure 1B). Second, we analyzed the structure of the resulting correlation network by forming “modules” (Figure 1C). Third, we performed an in-depth analysis of selected modules using probabilistic graphical models to identify a “network” of key biomarkers that represents the module (Figure 1D). Fourth, using the key biomarkers, we performed clustering of individuals to partition the study cohort into distinct health profiles with corresponding biomarker signatures (Figure 1E). We further characterized the clusters and examined disease risk using individuals’ personal history and, when available, longitudinal disease diagnosis data. We validated our main findings using an independent TwinsUK validation dataset derived from 1,083 females.

### Multimodal Correlations and Modules

We calculated correlations for each cross-modality pair of normalized features and selected a list of 11,537 statistically significant associations (with the FDR controlled at 5%) out of 427,415 total cross-modality comparisons (see Methods). A breakdown of the selected associations per pair of modalities is shown in Figure 2A. The largest number of significant associations (n=5,570) was observed between metabolome and clinical labs. This was previously observed by Price *et al.*^8^ and is mainly explained by separate measurements of the same or similar metabolites by the two modalities. The second largest number of significant associations (n=2,031) was between the metabolome and microbiome, given the large number of features measured in the two modalities (3% of the possible correlations between the two modalities were statistically significant; Figure 2A). We further discuss some of these associations in the subsequent sections. The third largest number of significant associations (n=1,858; 17%) was between metabolome and body composition. Some of the important findings from these two modalities have been discussed in Cirulli *et al.*^13^.

**Figure 2.**
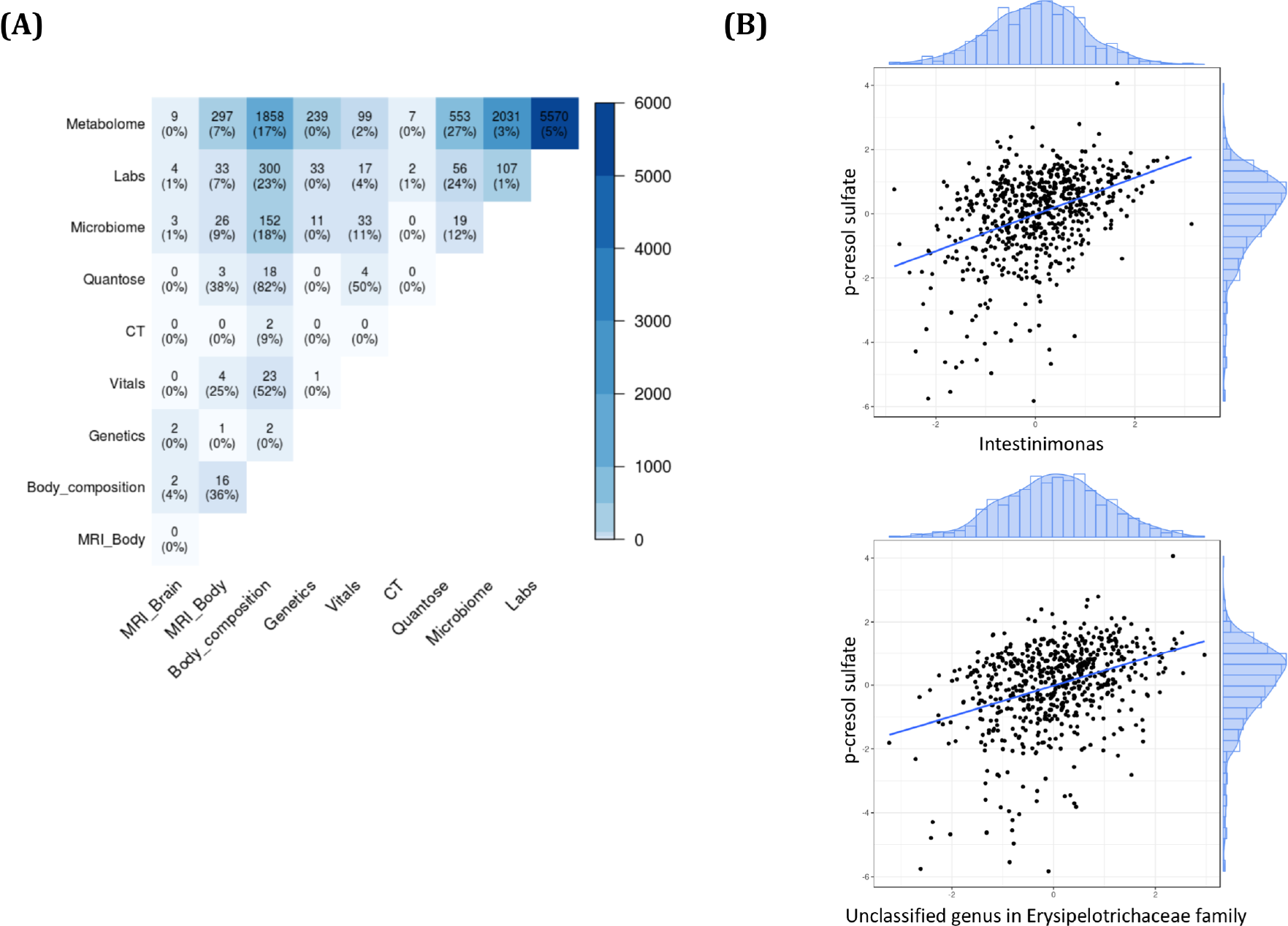
The number of significant cross-modality correlations for each pair of modalities is shown in part **(A)**. The percentages shown are the proportion of correlations that were significant out of all possible pairwise associations between the modality-pair. **(B)** Associations between p-cresol sulfate metabolite and (top) abundance of *Intestinimonas* genus, and (bottom) an abundance of unclassified genus in *Erysipelotrichaceae* family.

The most significant associations, apart from those between metabolome and labs, were expected correlations supporting well-established prior clinical research. Examples include associations between body mass index (BMI) and liver fat percentage (*p* = 1.35E-46) and between visceral adipose tissue (VAT) and insulin resistance (IR) score (p-value = 2.09E-44). These correlations highlight the importance of preventative medicine recommendations for reducing BMI and VAT, which are known risk factors for diabetes and other metabolic syndromes. We also observed height and polygenic risk scores (PRS) for height to be significantly correlated (*p* = 2.32E-44), highlighting the utility of genetics for trait prediction. Other significant genetic associations were observed between the PRS of lipid levels (high density lipoprotein, low density lipoprotein, total cholesterol, and triglyceride) and their corresponding lab measurements. Overall, we observed less than one percent of associations with genetic features that were significant. Conversely, body composition features (e.g. BMI, VAT, android/gynoid ratio, fat mass and lean mass) had the highest percentages of significant associations with several modalities (Figure 2A).

In addition, we observed novel associations between the metabolite *p*-cresol sulfate (pCS) and the microbiome genus *Intestinimonas* as well as an unclassified genus in the *Erysipelotrichaceae* family (*p* = 2.92E-24 and *p* = 2.98E-20 respectively; Figure 2B). pCS is a known microbial metabolite^14,15^ that is associated with accelerated cardiovascular disease and renal disease progression, and is considered a potential uremic toxin^14,16–18^. It is a sulfated phenolic compound generated in the colon by the bacterial fermentation of tyrosine. pCS was also associated with species diversity (*p* = 6.54E-19), and several genera (*Pseudoflavonifractor*, *Anaerotruncus*, *Subdoligranulum*, and *Ruminiclostridium*) in the *Ruminococcaceae* family (*p* = 9.52E-32, *p* = 1.39E-23, *p* = 9.48E-19, and *p* = 3.26E-11 respectively). Only the associations of pCS with species diversity and the *Ruminococcaceae* family have been observed previously^18–20^. These associations were validated in the independent TwinsUK cohort (see Methods; Table S2).

The significant associations from all modalities were then used to construct a network, which was used to find highly connected sets of variables that we refer to as *modules* (see Methods). Intuitively, the modules should group together markers that are biologically related. To avoid having the structure of the correlation network mostly determined by metabolome and clinical lab associations, we removed all of the corresponding edges. The result was two modules with, by far, the largest number of features (n>100 each) and numerous smaller modules.

Several of these modules permit biological interpretations. The largest of them was a *cardiometabolic module* containing many markers associated with cardiac disease and metabolic syndrome, as previously observed by Price *et al.*8. The second largest module was predominantly composed of microbiome taxa abundance, and metabolites that are known to be biomarkers for diversity in the gut microbiome. We refer to it as the *microbiome richness module*. Next, we present further detailed analysis on these two largest modules.

### Cardiometabolic Module

The largest module in the association network contained 355 nodes from clinical labs, metabolome, quantose, CT, microbiome, vitals, genetics, MRI-body and body composition data modalities. We ranked the features in this module by their relative centrality in the module using an eigenvector centrality score (see Methods), and we verified the presence of several markers associated with obesity, heart disease, and metabolic syndrome. The most central features for the module were VAT, BMI, liver fat percentage, lean mass percentile, glucose levels, blood pressure, triglycerides levels, IR score, several lipid metabolites, and several microbiome genera, including butyrate-producing bacterial genera such as *Pseudoflavonifractor*, *Butyrivibrio*, *Intestinimonas*, and *Faecalibacterium*.

#### Network Analysis for Key Biomarker Selection

While the module provides a general overview of how these features are interconnected, its construction is based only on pairwise associations. As such, it contains a significant amount of redundancy (e.g., two metabolites from the same pathway are likely to be connected to the same features from other modalities) and transitive edges (i.e., if and A and B are associated, and B and C are associated, we are likely to observe an association between A and C). In order to obtain a more meaningful representation of the interaction between the features in the module, we picked the 50 most central features and computed the inverse covariance matrix. This matrix defines a new network (called the *Markov network*) on these 50 features with the property that features A and B are only connected if they are correlated conditioned on all other features. The resulting network is shown in Figure 3A.

**Figure 3.**
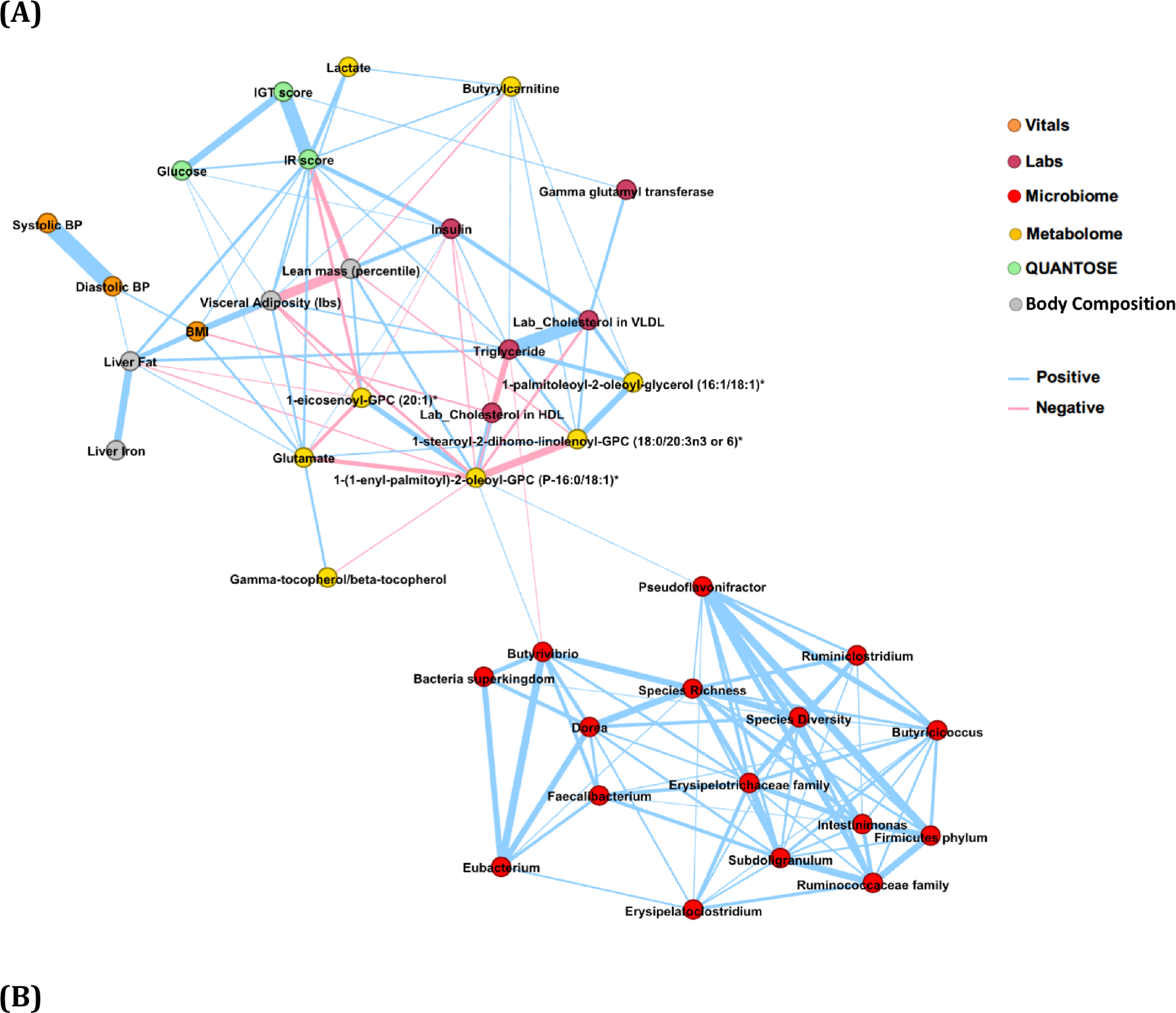
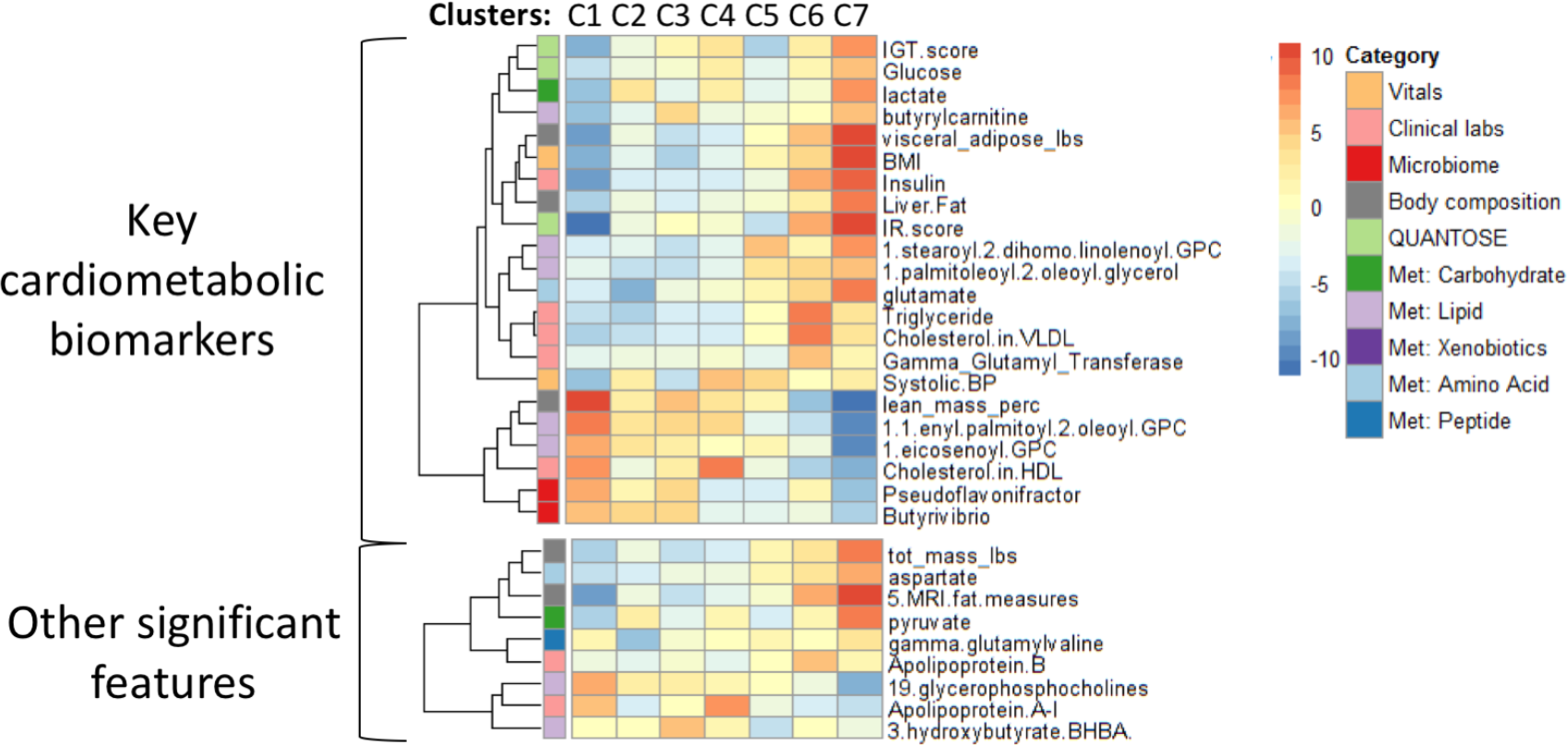
The cardiometabolic module. **(A)** We built a Markov network to identify the key biomarker features that represent the cardiometabolic module. This network highlights the most important associations after removing edges corresponding to indirect associations. We observed that the microbiome genera *Butyrivibrio* and *Pseudoflavonifractor* are the most relevant microbiome genera in the context of this module that interface with features from other modalities. **(B)** We clustered individuals using the key biomarkers. The heatmap shows z-statistics from logistic regression for an association between each cluster and each feature. The plot on the left shows the 22 key cardiometabolic biomarkers. The plot on the right shows associations that emerged from an analysis against the full set of 1,385 features with *p* < 1×10^-10^ as well as 3-hydroxybutyrate (BHBA) and Apolipoprotein B because of their particular enrichment in Clusters 3 and 6, respectively. Some correlated features have been collapsed, with the mean z-statistics displayed; the full set of features can be found in Figure S1. All of these significant associations showed consistent directions of effect in the TwinsUK cohort (details in Table S3); however, the microbiome features and 5 of the glycerophosphocholines were not measured in the TwinsUK cohort and thus could not be assessed for replication. Met = metabolome.

The Markov network emphasizes the most direct connections in the module. It suggests that (a) microbiome genera *Butyrivibrio* and *Pseudoflavonifractor* are “closest” to the remainder of the cardiometabolic module via a lipid metabolite 1-(1-enyl-palmitoyl)-2-oleoyl-GPC (P-16:0/18:1) and serum triglyceride, (b) systolic and diastolic blood pressure are mostly redundant from the point of the central variables in the module, demonstrated by the thickness of the edges, and (c) liver iron and gamma-tocopherol/beta-tocopherol are only associated to the rest of the module through other variables in their respective modalities. These observations allow us to determine a pruned set of 22 key cardiometabolic features (referred to as key biomarkers).

The key biomarkers included established features for cardiac and metabolic conditions (such as BMI, blood pressure, glucose levels and HDL) but also novel biomarkers (such as several metabolites and microbiome genera) that distinguish susceptibility to disease morbidity (Figure 3A). High abundance of the microbiome genera *Butyrivibrio* and *Pseudoflavonifractor* were well correlated with “good” cardiometabolic health (defined using traditional markers such as BMI, blood pressure and lipid levels). Several metabolites stood out as markers for the healthy profiles such as 1-(1-enyl-palmitoyl)-2-oleoyl-glycero-3-phosphocholine (GPC) and 1-eicosenoyl-GPC, and for the unhealthy profiles such as glutamate, butyrylcarnitine, lactate, 1-stearoyl-2-dihomo-linolenoyl-GPC, and 1-palmitoleoyl-2-oleoyl-glycerol.

#### Clustering of Individuals and Characterization

To assess the relationship between the health status of individuals and these 22 key biomarkers, we stratified individuals using hierarchical clustering. This clustering resulted in seven groups, each with a unique biomarker profile (Figure 3B). To better characterize the clusters, we compared each cluster to the full set of 1,385 features. We identified 106 features beyond the 22 used to calculate the cardiometabolic clusters that were significantly (*p* < 5.1E-06) enriched in at least one cluster compared to the others (Figures 3B and S3, and Table S3). Of the 78 features that were also measured in our validation cohort (TwinsUK baseline), 97.8% of the associations had consistent directions of effect in our validation cohort, and 77.8% were statistically significant (replication *p* < 3.9E-04; Table S3).

Based on BMI alone, clusters 1-4 can be roughly thought of as the healthy clusters and 5-7 as the unhealthy clusters. The individuals in cluster 1 can be characterized as containing the healthiest individuals, with a markedly higher lean mass percentile and low IR score. This cluster is also notable for its lower blood pressure, lower butyrylcarnitine levels, and higher HDL. The IR score and lean mass percentile for cluster 2 and 3 indicated individuals who were not as fit and healthy as those of cluster 1. In addition, cluster 2 displays the lowest glutamate values, while cluster 3 is characterized by the lowest blood pressure and the highest levels of 3-hydroxybutyrate. Cluster 4 is distinguished by an Impaired Glucose Tolerance (IGT) score that is higher than in the other clusters with healthy individuals and high levels of Apolipoprotein-A (Apo-A) and HDL cholesterol. Cluster 5 contains largely overweight individuals who nonetheless have low IR scores and low IGT. Cluster 6 contains mostly overweight and obese individuals with high android/gynoid ratios and IR scores; the individuals in this cluster were specifically characterized by the highest Apo-B, very low-density lipoprotein cholesterol and triglycerides of any cluster. Cluster 7 contains the least healthy individuals with respect to the markers under consideration, with a high prevalence of obesity, body fat and insulin resistance.

#### Disease Prevalence in Clusters

In addition to associations with features, we also compared rates of previously diagnosed cardio-metabolic conditions (i.e., diabetes, hypertension, hypercholesterolemia, heart disease, and stroke) between the clusters. We found significant differences between clusters in their rates of diabetes and hypertension diagnoses (Fisher’s exact *p* = 1.0E-04 and 2.3E-04, respectively). The findings were confirmed in the validation cohort (Fisher’s exact *p* = 4.3E-04 and < 1.0E-06, respectively) (Figure 4). Specifically, cluster 7 had significantly higher rates of diabetes, while cluster 1 had significantly lower rates of diabetes and hypertension. There were no significant differences between the clusters in heart disease or stroke history, though hypercholesterolemia showed a trend toward group differences in both cohorts that requires further validation (*p* = 0.03 in the discovery cohort; *p* = 0.09 in the validation cohort).

**Figure 4.**
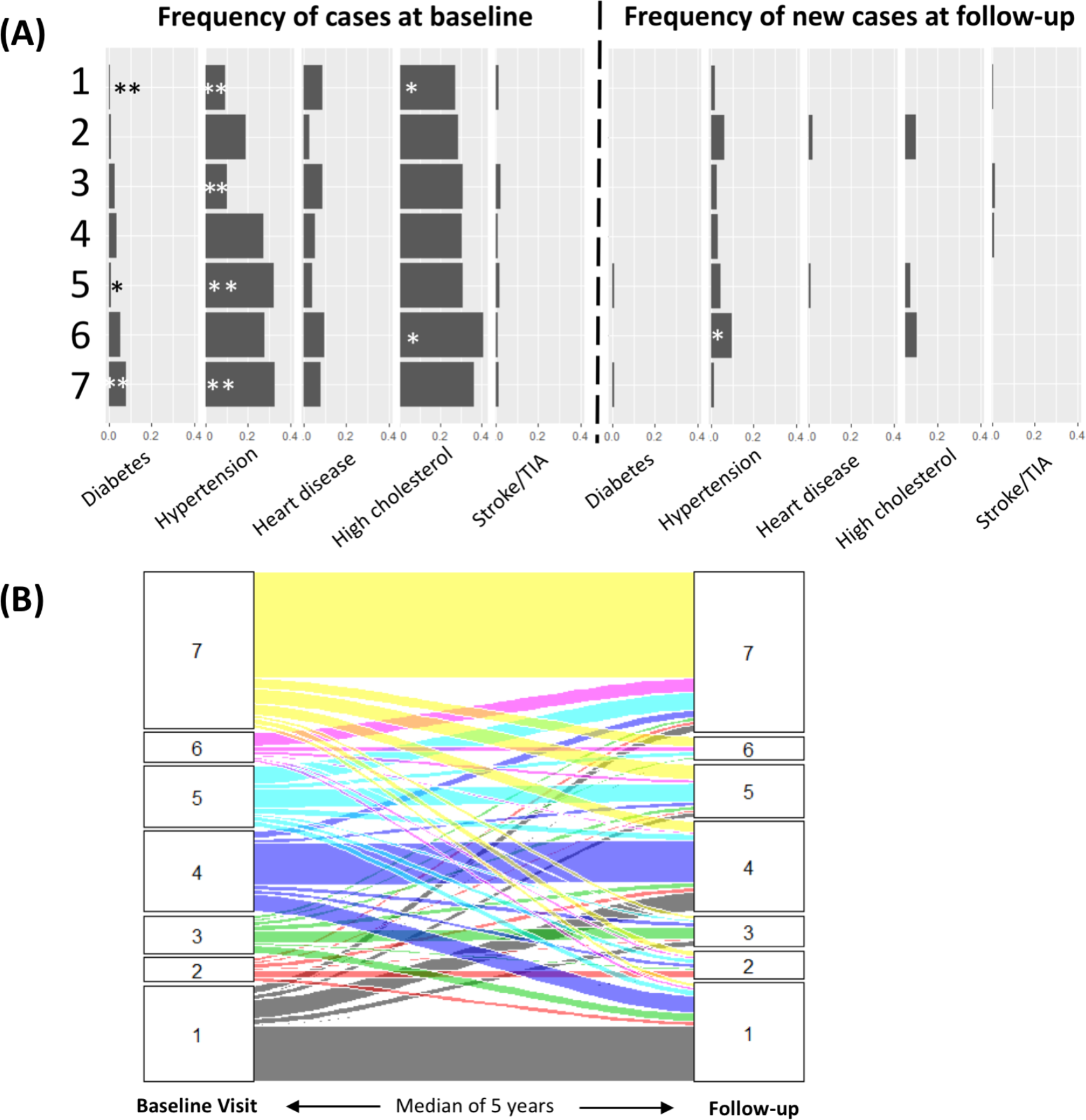
Disease enrichment and longitudinal outcomes of cardiometabolic clusters. **(A)** Bar plots showing the prevalence of disease at baseline (combined discovery and TwinsUK baseline cohorts; Figure S2 shows them individually) and the incidence of disease (i.e., only the new cases of disease) after a median of 5.6 years of follow-up (TwinsUK cohort). For Fisher’s exact test comparison of the rate in each cluster vs. the other clusters, \**p*<0.05, \*\**p*<0.005. **(B)** The rates at which individuals from each cluster transition into other clusters after a median of 5.6 years of follow-up. The plot shows individuals per cluster (1 to 7) at baseline visit that transition to other clusters during the follow-up. TIA = Transient Ischemic Attack.

Interestingly, cluster membership was a better predictor of diabetes diagnoses than were the traditional clinical features used to determine diabetes status: glucose, IGT score, and IR score, as well as BMI. Even after accounting for these traditionally predictive features, individuals in cluster 7 were significantly more likely to have diabetes than were members of the other clusters (logistic regression *p* = 0.01). The enrichment of hypertension diagnosis in cluster 7, however, was explained by blood pressure measurement, as expected.

The cardiometabolic key biomarkers that were the largest drivers of this association between diabetes and cluster 7 were the IR score, percent lean body mass, and the metabolites 1-stearoyl-2-dihomo-linolenoyl-GPC (18:0/20:3n3 or 6) and 1-(1-enyl-palmitoyl)-2-oleoyl-GPC (P-16:0/18:1). In a multivariable logistic regression containing the 22 cardiometabolic key biomarkers, the above mentioned were the four features that were significantly associated with diabetes status. They were also significant predictors of cluster 7, in addition to liver fat, HDL cholesterol, *Pseudoflavonifractor*, and the metabolites lactate and 1-eicosenoyl-GPC (20:1)).

We next sought to identify features that distinguished those in cluster 7 who did and did not have diabetes. We compared these two groups within cluster 7 for all 1,385 features, and the metabolite citrulline emerged as by far the best predictor of diabetes status, with decreased levels of citrulline being found in diabetes patients (*p* = 5.5E-06). We specifically found that this signal was due to diabetic patients on metformin medication, as opposed to other medications, a result consistent with studies showing decrease in serum concentration of citrulline after metformin therapy^21,22^.

We additionally investigated whether individuals with known rare pathogenic variants associated with obesity, cardiovascular conditions, or diabetes were enriched in any of the clusters (see Methods). However, we only found 3 individuals with known pathogenic/likely pathogenic variants: a pathogenic *TTR* variant (rs76992529; p.Val142Ile) for cardiomyopathy in a cluster 7 individual, a pathogenic *WFS1* variant (rs71530923; p.Arg42Ter) for Diabetes with insipidus, optic atrophy, and deafness in a cluster 1 individual, and a likely pathogenic *HNF1A* variant (rs193922587; p.Leu555Phe) for maturity onset diabetes of the young in a cluster 1 individual.

#### Longitudinal Disease Outcome

Our validation cohort was followed for a median of 5.6 (range 1.2-10.1) years, providing us with the opportunity to examine the longitudinal health trends in each cluster. During this follow-up, we observed 2 new diagnoses of diabetes, 2 cardiovascular events (angina and myocardial infarction), 7 strokes or transient ischemic attack (TIA), 24 new cases of hypertension, and 37 new cases of hypercholesterolemia. We found a significant difference between clusters in the number of new hypertension cases (Fisher’s exact *p* = 0.009). Specifically, those in cluster 6 were at higher risk for developing hypertension, and this association remained significant after controlling for baseline blood pressure, BMI, and age (logistic regression *p* = 0.002).

We also examined cluster membership after the follow-up (Figure 4). We found that cluster membership was fairly stable longitudinally, with 51.1% of individuals staying in the same cluster at the follow-up visit. For each cluster except cluster 6, the most common outcome at the follow-up visit was to remain in the same cluster. Cluster 6 had a very different pattern, with 84.3% of its members transitioning to other clusters, of which 55.8% moved to cluster 7. As cluster 7 is the least healthy cluster in terms of obesity, hypertension, and diabetes, this propensity of cluster 6 individuals to transition into cluster 7 individuals over time supports the idea of cluster 6 membership as an early precursor to a poor health outcome. Indeed, rates of hypertension were not significantly enriched in cluster 6 in the TwinsUK or combined cohorts at baseline but were after follow-up (though there was a trend for more hypertension cases in cluster 6 at baseline in our discovery cohort, *p* = 0.02). Our analysis therefore supports the classification of cluster 6 individuals as being at risk and prioritized for intervention before they progress to being truly unhealthy.

### Microbiome Richness Module

The microbiome richness module in the association network contained 167 features, the majority of which were from the metabolome (n=98) and microbiome (n=49) modalities. Similar to the in-depth analysis for the cardiometabolic module, we performed a network analysis to identify key biomarkers of this module and group individuals into clusters to assess their health status. Since microbiome was only measured for the last visit in our longitudinal validation cohort, we were unable to perform longitudinal disease outcome analysis for this module.

#### Network Analysis for Key Biomarker Selection

We selected the 40 most central nodes to construct a Markov network after ranking the nodes in the microbiome richness module by eigenvector centrality. The resulting network is shown in Figure 5A. The Markov network identifies the interface between the microbiome taxa and the metabolites in this module. In particular, we observed that most of the associations between the microbiome and the metabolome were mediated by species richness (i.e. the number of species present at a relative abundance greater than 10^-4^). Hence, we refer to this module as the microbiome richness module.

**Figure 5.**
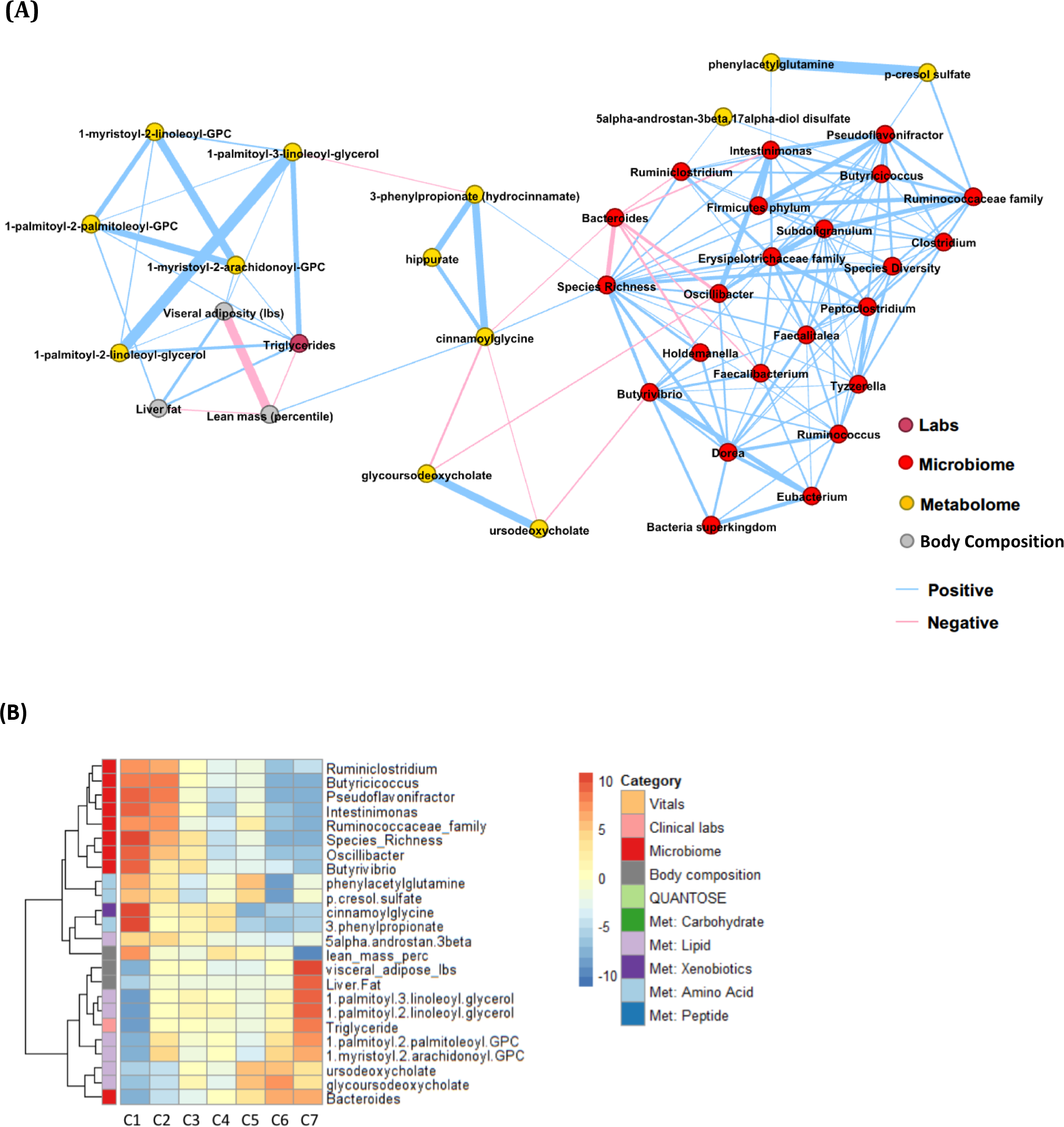
The microbiome richness module. **(A)** We built a Markov network to identify the key biomarker features that represent the microbiome richness module. Most of the associations between the microbiome and the metabolome were mediated by species richness. **(B)** We clustered individuals using the key biomarkers. The heatmap shows z-statistics from logistic regression for an association between each cluster and each feature. The plot on the left shows the 24 key biomarkers representing the module. Met = metabolome.

Specifically, species richness is connected to the metabolome via the mutually connected cinnamoylglycine, hippurate, and 3-phenylpropionate. This relationship is in agreement with a previous study15 that showed cinnamoylglycine and hippuric acid were not found in germ-free mice, and that 3-phenylpropionic acid is a metabolic product of anaerobic bacteria. Furthermore, a recent study^19^ identified hippurate, pCS, and 3-phenylpropionate as metabolic markers for microbiome diversity, with hippurate being the strongest of the three. Our approach suggests that cinnamoylglycine, in addition, is a good metabolic marker for gut microbiome health and that species richness is more directly associated with these metabolic markers than species diversity.

#### Clustering of Individuals and Characterization

As in the case of the cardiometabolic module, we selected our key biomarkers by excluding features that were only connected to their own modality in the Markov network. This resulted in 24 key biomarkers, which we then used to group individuals into 7 clusters (Figure 5B). The lipid profile that characterized this module had the lowest levels in cluster 1 and the highest levels in cluster 7. While, the microbiome genera abundances and species diversity were the highest in cluster 1 and the lowest in 7. The exception was *Bacteroides*, which showed the opposite trend. Associations with the full set of 1,345 features showed that cluster 7 could be characterized as the least healthy, with the highest levels of body fat, BMI, triglycerides and total cholesterol and the lowest lean mass. Cluster 1 had values at the opposite extreme for each of these traits and can be characterized as the healthiest. In addition, the clusters were largely distinguished by differences in various lipids and microbiome genera. The enrichments most specific to each cluster were as follows: Cluster 1 had the highest levels of hippurate and cinnamoylglycine; Cluster 2 had the highest levels of *Holdemania*, *Oscillospiraceae*, *Ruminococcaceae*, and *Alistipes*; Cluster 3 had the lowest levels of *Flavonifractor* and 3-methylglutaconate; Cluster 4 had the lowest levels of *Lachnospiraceae*; Cluster 5 had the highest levels of 4-hydroxyhippurate and 3’-3-hydroxyphenyl propionate and the lowest levels of cinnamoylglycine; Cluster 6 had the highest levels of glycoursodeoxycholate and lowest levels of p-cresol sulfate, p-cresol glucuronide, phenylacetylglutamine, 6-hydroxyindole sulfate and *Ruminiclostridium*; and Cluster 7 had the lowest levels of HDL and the highest urate, ferritin, Apo-B, and diastolic blood pressure levels.

While the clusters seemed to reflect different states of gut microbiome health, which may be associated with overall cardiometabolic health, we found no enrichment of cardiometabolic or other diseases in any of the clusters.

### Comparing Membership Across the Modules

We proceeded to compare the membership of individuals in the clusters from the cardiometabolic and the microbiome richness modules. There was significantly (*p* < 0.001) more overlap of individuals between clusters 7 in the two modules and also between clusters 1 than expected by chance: 66% of those in the microbiome richness cluster 7 were in the cardiometabolic cluster 7, and 45% of those in the microbiome richness cluster 1 were also in the cardiometabolic cluster 1. In contrast, only 1% of those in microbiome richness cluster 7 were in cardiovascular cluster 1 (Figure S4).

## Discussion

We analyzed 1,385 multi-modal features collected from 1,253 individuals using an ensemble of unsupervised machine learning and statistical approaches. We identified novel associations and novel biomarker signatures that stratified individuals into distinct disease states including early disease transition states. The main findings were replicated in an independent validation cohort of 1,083 females (TwinsUK).

Specifically, we performed association analysis of features across modalities and found novel significant associations between *p*-cresol sulfate (pCS) and the microbiome genera *Intestinimonas* and an unclassified genus in the *Erysipelotrichaceae* family. pCS is a known microbial metabolite and is considered to be an uremic toxin. It is produced by bacteria fermenting undigested dietary proteins that escape absorption in the small bowel^23–25^. It appears to be elevated in the sera of chronic kidney disease (CKD) patients, and it is associated with increased mortality in patients with CKD26 and an increased risk of cardiovascular events^26^. The *Intestinimonas* genus is known for being a butyrate-producing species that digests lysine and fructoselysine in the human gut^27^, but it is otherwise not well described. Members of the *Erysipelotrichaceae* family might be immunogenic and can potentially flourish after treatment with broad spectrum antibiotics^28^. An increased abundance of *Erysipelotrichaceae* has been observed in obese individuals, and several other lines of evidence suggest a role in lipid metabolism^28^. Our novel associations were validated in the TwinsUK cohort and could further be analyzed as potential therapeutic targets to decrease pCS levels and its toxicity.

Community detection analysis of the 11,537 statistically significant feature associations identified two primary modules of densely connected features: the cardiometabolic module and the microbiome richness module. Both of these modules identified individuals who were healthier, according to measures like their BMI, and metabolic and cardiovascular disease status, and individuals who were less healthy. Interestingly, when grouping individuals with distinct signatures in each module together into clusters, the healthiest group of the cardiometabolic module largely overlapped the healthiest group of the microbiome richness cluster. The same was observed for the least healthy (i.e. “unhealthiest”) cluster. Such co-enrichment of individuals in the unhealthiest clusters derived from both modules suggests patterns of comorbidity and highlights the interaction between cardiometabolic health and gut microbiome health.

The key biomarkers identified in the cardiometabolic module consisted of potentially novel features in addition to the traditional clinical features from several modalities. The potentially novel biomarkers included the abundance of the microbiome genera *Butyrivibrio* and *Pseudoflavonifractor* and several metabolites, such as 1-(1-enyl-palmitoyl)-2-oleoyl-GPC, 1-eicosenoyl-GPC, glutamate, and 1-stearoyl-2-dihomo-linolenoyl-GPC. The higher abundance of the two microbiome genera has been associated with decreased adiposity and improved insulin sensitivity. The *Butyrivibrio* genus is known for its butyrate-producing species and plays a major role in fiber and other complex polysaccharide degradation^29,30^. An increased abundance of *Butyrivibrio* increases the rate of butyrate production, which is suggested to decrease risk of type 2 diabetes and decrease adiposity^31,33,35^. In addition, the oral administration of a *Butyrivibrio* species was shown to reduce putative preneoplastic lesions in mice, suggesting a role for the microbiome species as a probiotic in the prevention or suppression of colorectal cancer^31^. A weight-loss study showed enrichment of *Pseudoflavonifractor* at baseline in individuals who succeeded in losing their weight consistently for two years^37^. In our study, we observed a higher abundance of *Butyrivibrio* and *Pseudoflavonifractor* in individuals in cluster 1, which is consistent with our observation of a very low prevalence of diabetes, hypertension, and obesity in that cluster.

We identified another potential biomarker for health from the analysis of the microbiome richness module -- the metabolite cinnamoylglycine was associated with microbiome species richness and lean mass percentage. It was observed to be abundant in individuals in cluster 1, representing healthy individuals. Cinnamoylglycine is related to gut bacterial metabolism, and it was identified as being present only in the serum or colonic lumen from conventional but not germ-free mice15. Additional study is needed to confirm the role of cinnamoylglycine on health and to understand its biological mechanism.

We found that the cluster membership for individuals was a better predictor of diabetes than the traditional clinical biomarkers such as glucose, BMI and insulin resistance. The novel biomarkers in the diabetes signature included 1-stearoyl-2-dihomo-linolenoyl-GPC and 1-(1-enyl-palmitoyl)-2-oleoyl-GPC. These lipid metabolites are not well studied but are likely present in cell membranes and fat-carrying vehicles such as HDL. A study on a related metabolite 1-palmitoyl-2-oleoyl-sn-GPC (POPC) suggested a role in insulin resistance^39^; glucose uptake in skeletal muscle showed that a synthetic reconstituted discoidal HDL made with POPC produced insulin-like effects. Future work on these metabolites may prove them to be novel biomarkers for insulin resistance and diabetes.

A longitudinal disease outcome analysis in the follow-up TwinsUK data found an early disease signature for hypertension (membership in the cardiometabolic module cluster 6). We also observed that more than half of the individuals from cluster 6 transition to cluster 7 (the unhealthiest cluster) in the follow-up visit, suggesting that cluster 6 membership is an early indication of a poor health outcome. These signatures can be used to prioritize individuals for intervention.

We did not observe a substantial number of significant findings for the genetic features. This result is not unexpected given the relatively small sample size considered here compared to the large sample sizes required for finding statistically significant association in genetic studies. Additionally, the analyses focus on the main/strongest findings from unsupervised pattern detection, and an overwhelming signal from other functional measurements dampens signals from genetics. The types of associations with the largest effect sizes would be for rare variants and diseases, for which any population-based cohort like the one studied here would be underpowered. Finally, the PRS derived using common variants for certain traits could only explain a small fraction of the variance; therefore, we are underpowered to detect significant associations.

In recent years, several organizations have begun gathering cohorts with high throughput data from multiple modalities. The collection of such datasets from large cohorts is a necessary step in systems medicine to gain comprehensive insights into an individual’s health status and to understand complex disease mechanisms. A systematic and supervised approach to analyzing an individual’s deep phenotype data, as shown in our previous publication^4^, is important for precision medicine screening. However, it is also crucial to perform unsupervised multimodal data analyses, as described here, to sift through this wealth of information for novel findings of signatures of health and disease. These novel discoveries and the characterization of complex interactions allow us to transition towards personalized, preventative health risk assessments.

## Methods

### Data Collection and Data Features

For the study, we collected data from 1,253 self-assessed healthy individuals in our clinical research facility. We used several tools and techniques referred to as *modalities* to collect the data. The modalities included whole genome sequencing (WGS), microbiome sequencing, global metabolome, insulin resistance (IR as defined by Cobb *et al.*^12^) and glucose intolerance (IGT as defined by Cobb *et al.*^32^) laboratory developed tests (Quantose™), whole body and brain magnetic resonance imaging (MRI), dual-energy x-ray absorptiometry (DEXA), computed tomography (CT) scan, routine clinical laboratory tests, personal/family history of disease and medication, and vitals/anthropometric measurements. Data collection has been described in detail in our previous manuscript on the first 209 individuals enrolled in a precision medicine study^4^. In addition to the modalities described in the previous study, we have now included CT scan and microbiome sequencing^10^. Not all data was collected on all individuals. The number of individuals and the number of features per modality are summarized in Table S1.

We performed CT scans on individuals over the age of 35 years. Patients were scanned during a single breath-hold using a 64-slice GE Healthcare EVO Revolution scanner (GE Healthcare, Milwaukee, Wisconsin). Gated axial scans with 2.5 mm slice thickness were performed using a tube energy of 120 kVp and the tube current adjusted for individuals’ body mass index. Images were subsequently analyzed using an AW VolumeShare 7 workstation (GE Healthcare, Milwaukee, Wisconsin) and regions of coronary calcification were manually identified in order to compute Coronary Artery Calcium (CAC) Agatstan scores^34^. We used Multi-Ethnic Study of Atherosclerosis (MESA)^36^ reference CAC values to calculate the percentile of calcification for each individual matched for age, sex and ethnicity.

For microbiome sequencing, we performed whole genome sequencing on stool samples to analyze the microbial communities^10^. For this modality, the features included species richness, species diversity, the fraction of human DNA, *Proteobacteria*, and the abundance of 72 genera^10,38^. Microbiome species richness is defined as the number of species present at a relative abundance greater than 10^-4^. Microbiome species diversity is defined as the Shannon entropy of the taxon abundance vector^40^.

Whole-genome sequencing data was used to compute the following features: polygenic risk scores (PRS)^41^ for 51 diseases and traits, HLA-type^42^, 30 known Short Tandem Repeats (STR) disease loci^43^, and known rare pathogenic variants from ClinVar (Set-1 and Set-2 from Shah *et al.*^44^). We also computed ancestry using the method described by Telenti *et al.*^9^ from WGS data.

### Data Normalization

Several features were correlated with age, sex or ancestry. To remove this bias, we used multiple linear regression to identify the covariates among age, sex and ancestry (first four principal components), that were significantly associated with each feature at a *p* < 0.01 significance level. We then replaced the feature values with the residuals after regressing the associated covariates.

To address the non-Gaussian distributions of various features from several modalities, we utilized a rank-based inverse normal transformation^45^. We applied this transformation to all microbiome abundance data, as these features exhibit fairly non-Gaussian distributions, and to any other feature in which more than 40% of the samples had the same value.

### Constructing Multimodal Correlation Modules

We performed Spearman correlation analysis and calculated p-values for each cross-modality pair of features. The correlation was calculated only if at least 30 samples had data for the pair of features. We selected statistically significant association using the Benjamini-Hochberg^46^ approach by limiting the false discovery rate to 5%.

The significant associations were used to construct a network where each feature from each modality is a *node*, and the corresponding association p-value is an *edge* between two features from different modalities. The weight of an edge is defined as -log(*p*), where *p* is the p-value of the corresponding Spearman correlation. We then used the Louvain algorithm^47^ to perform community detection. This algorithm was chosen as it is more aggressive and removes more edges than the one utilized in Price *et al.*^8^. We refer to each of the “communities” computed by the algorithm as modules.

We allowed for nodes to belong to multiple modules. This was permitted only when a node was assigned to a module by the community detection algorithm but had more than 20 significant associations with another module (or more associations with another module than it had with its assigned module).

### Key Biomarker Selection and Conditional Independence Network Construction

To perform a deeper analysis of the cardiometabolic and microbiome richness modules, an initial list of candidate biomarkers was selected using eigenvector centrality. More precisely, for the subnetwork corresponding to each of these two modules, we ranked all the nodes according to their eigenvector centrality score. For the cardiometabolic module, the 50 most central features were selected and for the microbiome richness module, the 40 most central features were selected.

For each of the two modules, the central features were mean-imputed and converted to Gaussian distributions using the rank-based inverse normal transformation as described above. These features were then used to construct a sparse network using the GraphLasso method^48^. This method estimates the inverse covariance matrix of the selected features using a lasso penalty to induce sparsity. The resulting network is a conditional independence network (also known as a “Markov Network”) in the sense that the absence of an edge between two features implies that they are approximately conditionally independent given the remaining features in the Markov network. Unlike the cross-modality correlation network, here the connections between features from the same modality were allowed and are typically the strongest associations. This method tends to be less sensitive than the pairwise Spearman associations initially computed, and several weak cross-modality associations observed previously were not observed in the Markov network.

### Clustering Individuals into Distinct Health Profiles

The Markov network allowed us to identify features that were associated with the rest of the network through other features in its own modality. By removing such features, we obtained our final set of biomarkers, both for the cardiometabolic module and for the microbiome richness module.

These key biomarkers were then utilized to cluster the individuals. We first selected individuals for which the key biomarkers were available (668 individuals for the cardiometabolic module and 640 individuals in the microbiome diversity module). The resulting data matrix had each feature scaled to have zero mean and unit variance. The missing values were imputed using softImpute^49^.

We performed hierarchical clustering on the set of individuals based on complete linkage and a correlation distance metric, and extracted clusters from the dendrogram.

### Statistical associations between clusters and other traits

We compared the rates of disease diagnoses and medication use across the seven cardiometabolic and the seven microbiome richness clusters. A Fisher’s exact test was used with the fisher.test command in R (using a Monte Carlo simulated p-value with 1E6 replicates) to test for statistical significance after Bonferroni correction for multiple tests.

We also compared the individuals in each cluster to all individuals not in that cluster for each of the 1,354 features using a logistic regression with the glm command in R. There was thus a separate analysis performed for cluster 1 vs. everyone else, cluster 2 vs. everyone else, etc. Significant associations were those that survived Bonferroni correction for multiple tests.

### Validation Cohort

For validation of our findings, we utilized 1,083 individuals from a study cohort (referred to here as “TwinsUK”) of largely European ancestry female twins enrolled in the TwinsUK registry, a British national register of adult twins^50^. The cohort included data from WGS, metabolome, microbiome, DEXA, clinical blood laboratory tests, and personal history of disease and medication. The data from the modalities was collected from three longitudinal visits over the course of a median of 13 years. To capture a population with adequate sample sizes for the overlapping modalities used in the present study, we restricted our analysis to data from visit 2 (referred here as “baseline”) and visit 3 (referred here as “follow-up”). Microbiome samples were only collected at visit 3. To be included in the analysis, phenotyping measurements were required to be collected within 90 days of the metabolome draw for each visit, or within 6 months for microbiome. For the validation of metabolome and microbiome correlations, we used only one of the twins to avoid bias from relatedness, totaling 538 individuals. For the cardiometabolic module analysis, we imputed liver fat, Gamma-Glutamyl Transferase (GGT), IGT, IR and glucose using regularized linear regression with L1 penalty (R glmnet package).

## Supplemental Information

**Figure S1.**
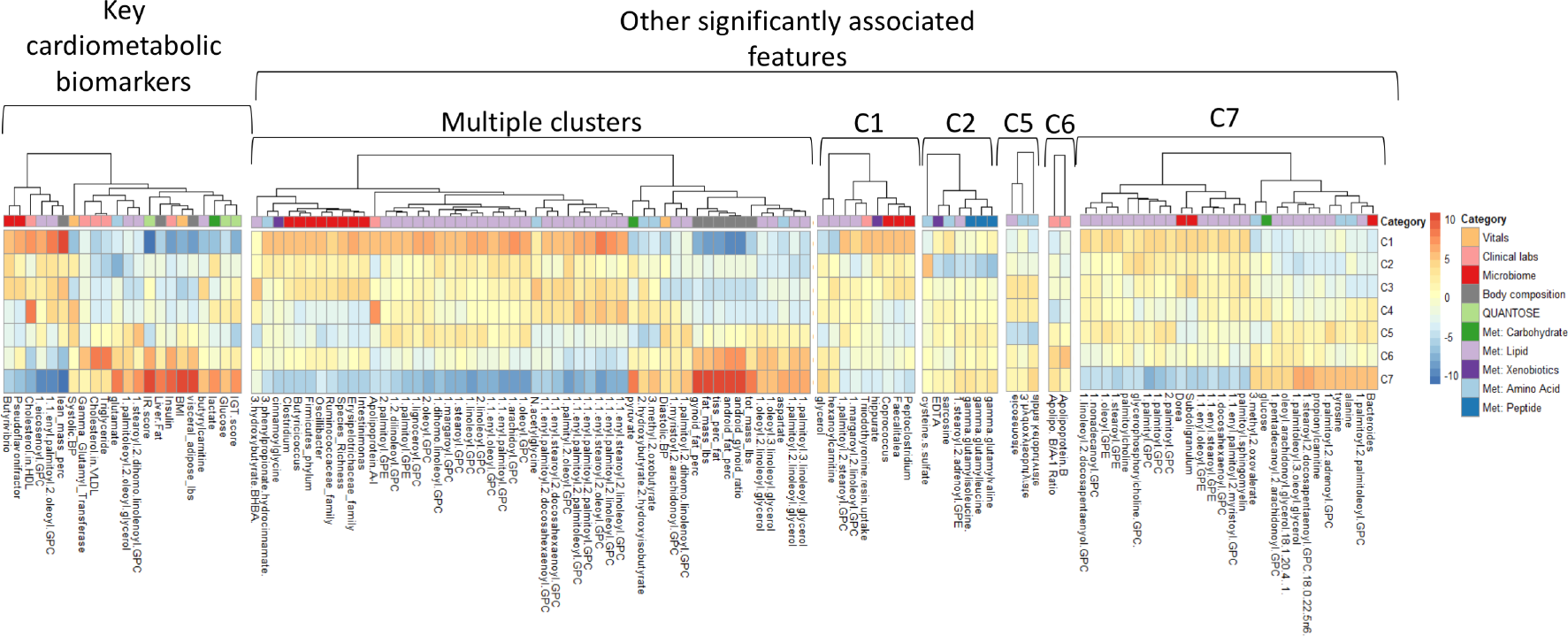
The cardiometabolic module: The heatmap shows the Z-statistics from a logistic regression for an association between each cluster and each feature. The plot on the left shows the 22 key cardiometabolic features. The plots on the right show significant associations that emerged from an analysis against the full set of 1,385 features. The first plot begins with the features that had significant associations with multiple clusters, and the remaining plots show features that were significantly associated with only one feature. The highlighted groups (e.g., lipid group 1) are largely internally redundant and were collapsed in Figure 4 by plotting their mean Z-statistics value. With the exception of 51 features that were not measured in the TwinsUK cohort (see Table S3), all associations showed directions of effect in the TwinsUK cohort that were consistent with the original association except for the associations between fat mass and visceral adipose and cluster 6, 1-stearoyl-2-docosapentaenoyl-GPC and cluster 7, and 3-methyl-2-oxobutyrate and cluster 5 (Table S3).

**Figure S2.**
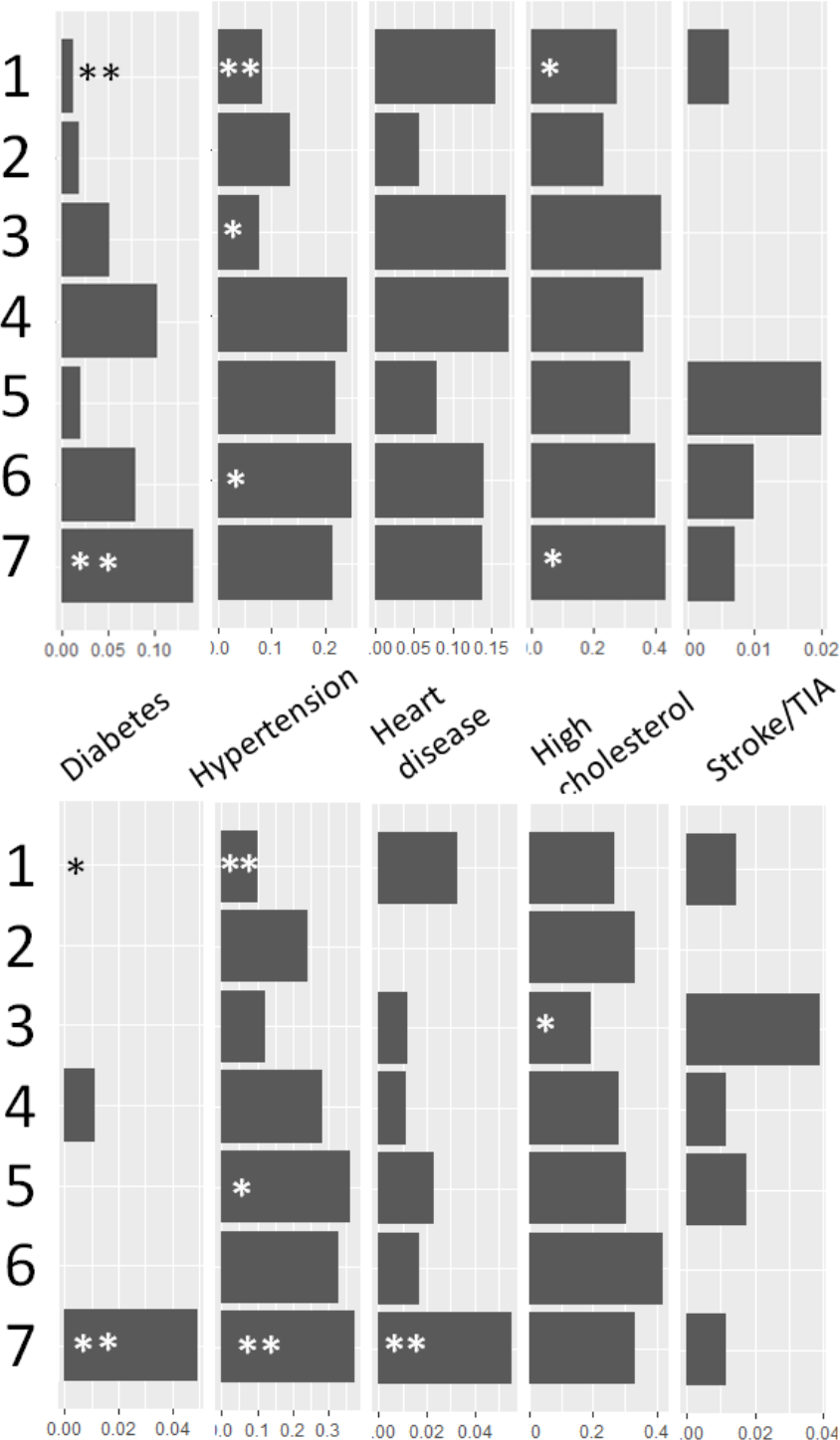
Prevalence of disease diagnoses. A) Discovery cohort, B) TwinsUK cohort at baseline. The combined cohort is shown in Figure 4A. For Fisher’s exact test comparison of the rate in each cluster vs. the other clusters, \**p*<0.05, \*\**p*<0.005. TIA = Transient Ischemic Attack.

**Figure S3.**
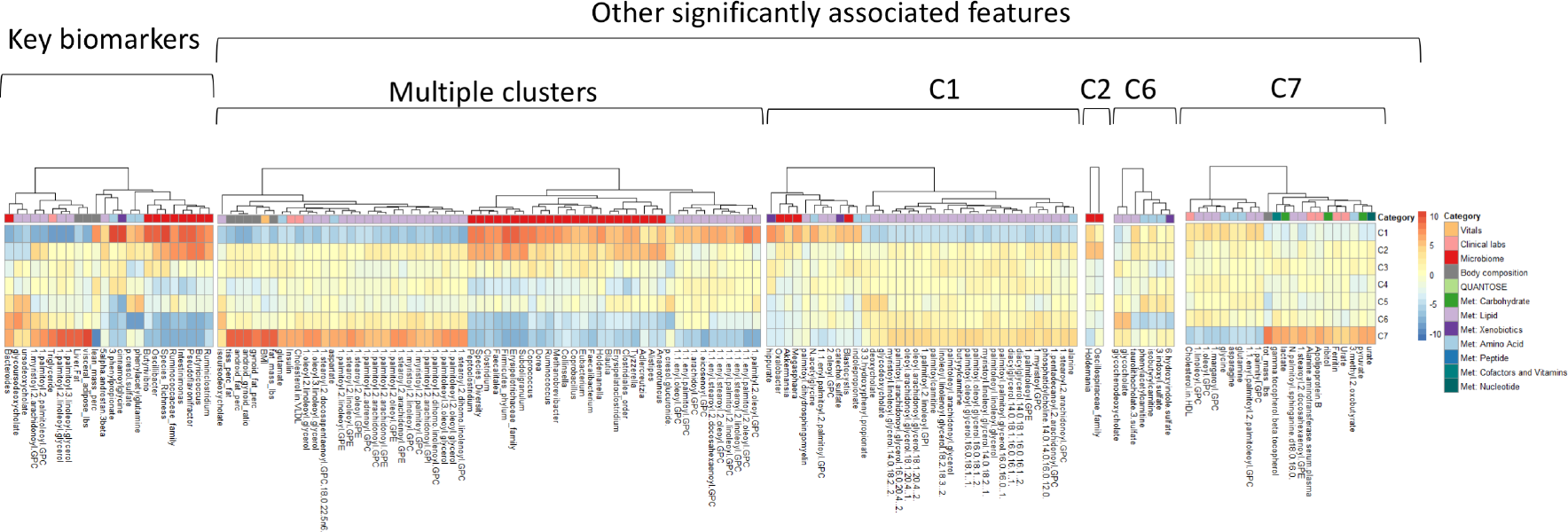
The microbiome richness module: The heatmap shows the Z-statistics from a logistic regression for an association between each cluster and each feature. The plot on the left shows the 24 key biomarkers. The plots on the right show significant associations that emerged from an analysis against the full set of 1,385 features. The first plot begins with the features that had significant associations with multiple clusters, and the remaining plots show features that were significantly associated with only one feature.

**Figure S4.**
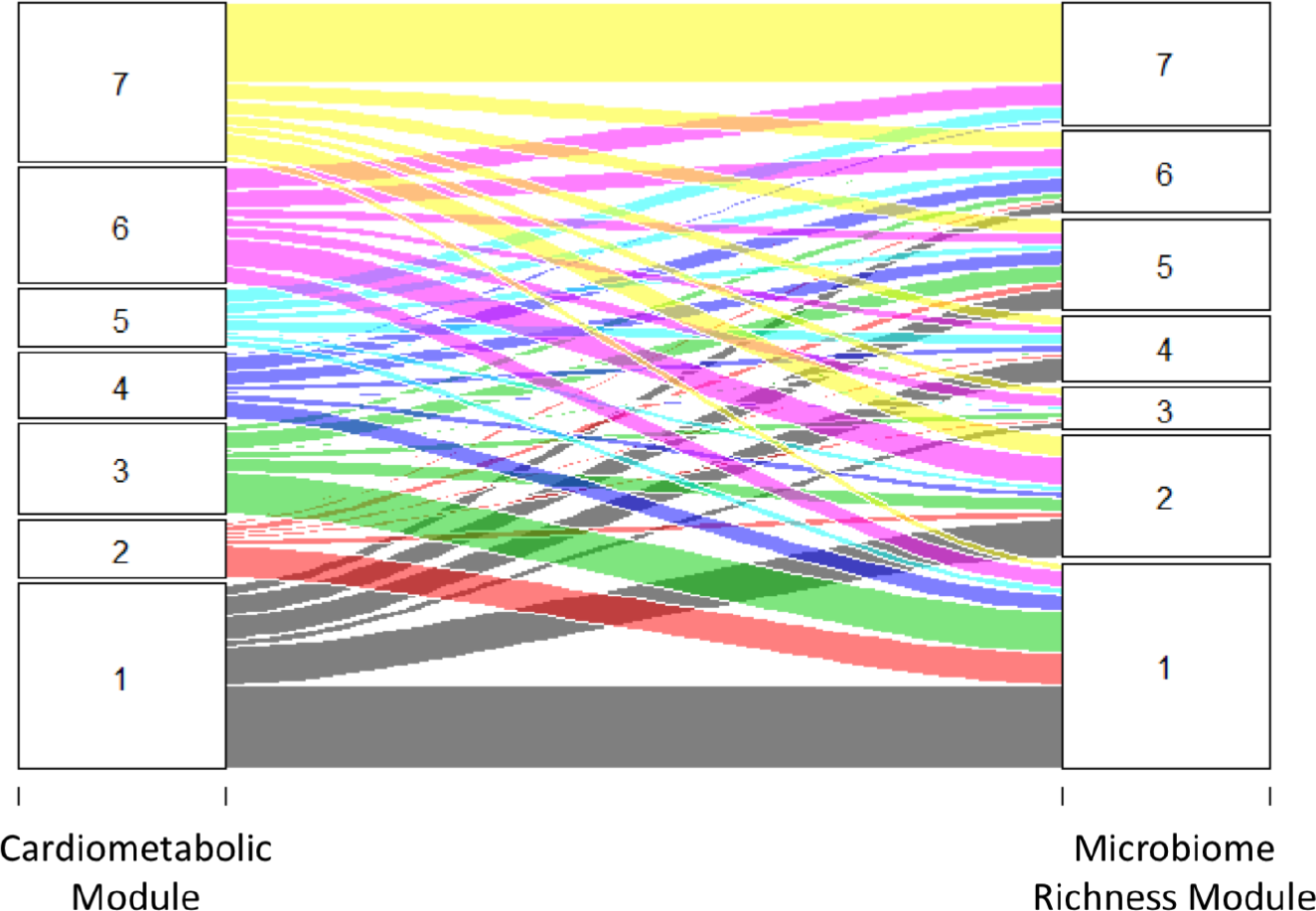
Overlap of clustered individuals between the cardiometabolic module and the microbiome richness module. There was a high overlap between the two clusters 1 and the two clusters 7. Of those in microbiome richness cluster 7, 66% were also in the cardiometabolic cluster 7, and 45% of individuals in cluster 1 of the microbiome richness module were also members of cluster 1 of the cardiometabolic module. In contrast, only 1% of those in microbiome richness cluster 7 were in cardiovascular cluster 1.

**Table S1.** The table shows number of individuals and number of features measured per modality. We combined body composition features from DEXA and MRI and treated them as a separate modality (“Body composition”). MRI = magnetic resonance imaging; DEXA = dual-energy X-ray absorptiometry; CT = Computed tomography.

| **Modalities** | **Number of individuals** | **Number of features** |
| --- | --- | --- |
| Genetics | 1253 | 182 |
| Personal/family medical history | 1253 | N/A |
| Vitals | 1253 | 4 |
| MRI | 1134 | 5 (Brain) + 4 (Body) |
| Microbiome | 877 | 76 |
| Clinical labs | 769 | 118 |
| Metabolome | 674 | 1010 |
| Quantose | 711 | 2 |
| Body composition | 605 (DEXA) + 1134 (MRI) | 11 |
| CT | 386 | 2 |

**Table S2.** The table shows microbiome genera that are associated with a metabolite *p*-cresol sulfate in both the discovery cohort and the replication cohort.

| **Microbiome Genus** | **Microbiome Family** | **Association P-value in discovery cohort** | **Association P-value in replication cohort** |
| --- | --- | --- | --- |
| Intestinimonas | Unclassified Clostridiales | 2.92E-24 | 8.54E-12 |
| Unclassified genus in Erysipelotrichaceae family | Erysipelotrichaceae | 2.98E-20 | 3.42E-04 |
| Pseudoflavonifractor | Ruminococcaceae | 9.52E-32 | 1.04E-09 |
| Anaerotruncus | Ruminococcaceae | 1.39E-23 | 1.85E-07 |
| Subdoligranulum | Ruminococcaceae | 9.49E-19 | 2.20E-06 |
| Unclassified genus in Ruminococcaceae family | Ruminococcaceae | 1.95E-38 | 1.20E-14 |
| Ruminiclostridium | Ruminococcaceae | 3.26E-11 | 6.06E-06 |

Table S3 The table shows features associated with the seven clusters of individuals using the cardiometabolic biomarker signature in both the discovery cohort and the validation cohort.

